# A predictive model of antibody binding in the presence of IgG-interacting bacterial surface proteins

**DOI:** 10.1101/2020.10.20.347781

**Authors:** Vibha Kumra Ahnlide, Therese de Neergaard, Martin Sundwall, Tobias Ambjörnsson, Pontus Nordenfelt

## Abstract

Many bacteria can interfere with how antibodies bind to their surfaces. This bacterial antibody targeting makes it challenging to predict the immunological function of bacteria-associated antibodies. The M and M-like proteins of group A streptococci exhibit IgGFc-binding regions, which they use to reverse IgG binding orientation depending on the host environment. Unraveling the mechanism behind these binding characteristics may identify conditions under which bound IgG can drive an efficient immune response. Here, we have developed a biophysical model for describing these complex protein-antibody interactions. We show how the model can be used as a tool for studying various IgG samples’ behaviour by performing in silico simulations and correlating this data with experimental measurements. Besides its use for mechanistic understanding, this model could potentially be used as a tool to predict the effect, as well as aid in the development of antibody treatments. We illustrate this by simulating how IgG binding in serum is altered as specified amounts of monoclonal or pooled IgG is added. Phagocytosis experiments link this altered antibody binding to a physiological function and demonstrate that it is possible to predict the effect of an IgG treatment with our model. Our study gives a mechanistic understanding of bacterial antibody targeting and provides a tool for predicting the effect of antibody treatments.

## Introduction

Many bacteria can interfere with how antibodies bind to their surfaces. Opsonisation, the process in which antibodies target pathogens through binding to facilitate phagocytosis, is essential for the defense against infections. However, several significant bacterial pathogens express surface proteins that interfere with this process by binding antibodies outside the antigen-binding region. A common target is the Fragment crystallizable (Fc) region on Immunoglobulin G (IgG) and proteins expressing IgGFc-binding regions include protein G from group C and G streptococci (2), protein A from *Staphylococcus aureus* (10) and M and M-like proteins from group A streptococci (GAS) (11) (29) (30). By binding the Fc region on IgG the bacteria can reduce the ability of the immune response to detect and eliminate the bacteria from the host organism through Fc-mediated phagocytosis (9) (14). This makes it difficult to predict the immunological effect of bacteria-associated antibodies.

According to our previous findings (22), the IgGFc-binding of M and M-like proteins is more prominent in low IgG concentration environments, corresponding to that of the human throat milieu. At the higher IgG concentrations in plasma, IgG is mostly bound to the surface proteins via Fab. This suggests that the antibody binding orientation is dependent on the IgG concentration. The mechanism for this behaviour is however unclear.

*Streptococcus pyogenes* causes more than 700 million uncomplicated throat and skin infections annually and can be at the root of rare but very serious invasive infections such as sepsis (4). It is estimated that group A streptococcal infections lead to more than 0.5 million deaths annually (4). Monoclonal antibody treatments have shown potential for a wide spectrum of disease. Despite this, there are very few monoclonal antibodies (mAbs) clinically available for treatment of bacterial infections (20) (19). Furthermore, pooled human IgG is used clinically for treatment of certain invasive GAS infections, but studies show no clear effect of this treatment (13) (15) (5) (3). A better understanding of the binding characteristics of M protein can help identify conditions under which bound IgG can drive an efficient immune response.

While there exist models of antibody binding to antigens that build on receptor-ligand kinetics and structural data (24) (23) (12), there is no model that describes the complex interaction of multiple antibody clones with varying affinities and multi-valency. Moreover, no model takes Fc binding into account. The binding of IgG to M protein in the host environment depends on total concentration of IgG, concentration of the different IgG clones, concentration of bacterial surface proteins, number of Fab-binding sites, location of epitopes and binding energies to these locations (**Fig. 1a**).

**Fig. 1.**
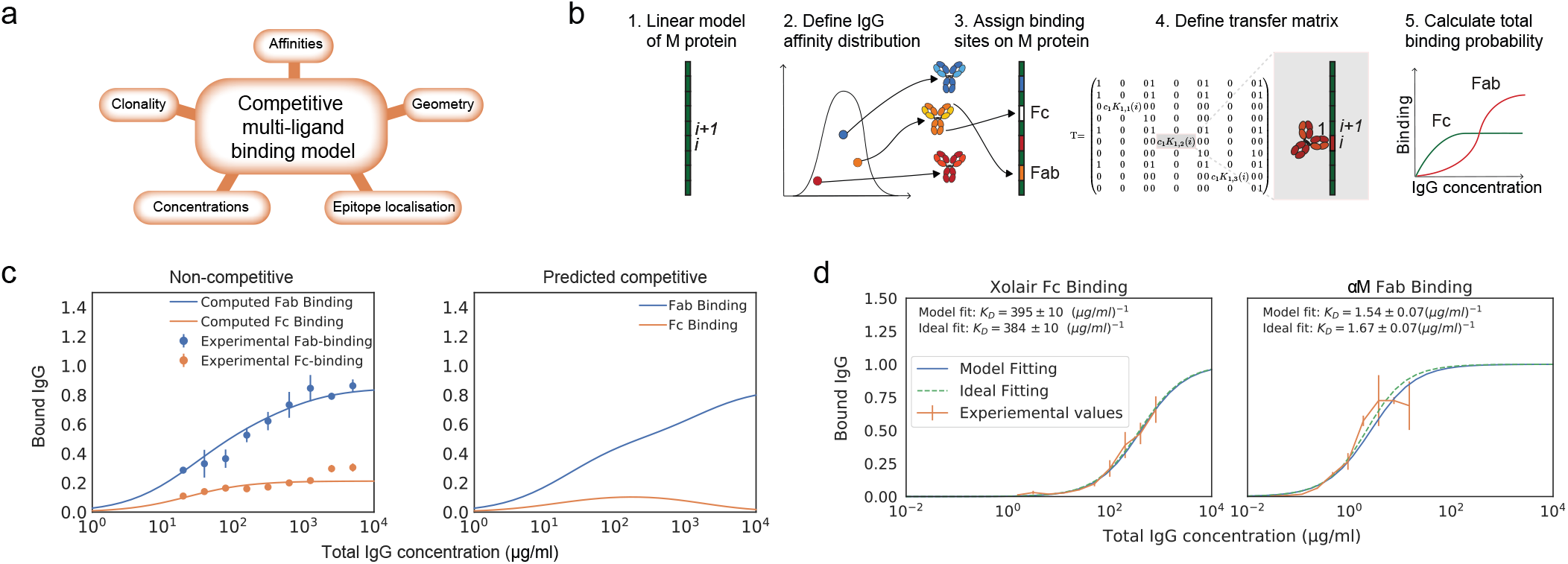
Biophysical modeling of competitive antibody binding to M protein. IgG binding orientation on bacterial surface proteins can be described through theoretical modelling. **a. Key parameters of the competitive polyclonal antibody binding model.** The binding of IgG to M protein is complex as it depends on the total concentration of IgG, concentration of the different IgG clones, the concentration of bacterial surface proteins, number of Fab-binding sites, location of epitopes and binding energies to these locations. These factors are taken into consideration in our biophysical model. **b. Framework for the competitive polyclonal antibody binding model.** 1. A linear-protein approximation of M protein is generated. 2. A polyclonal IgG sample can be modeled as a distribution of antibodies with different affinities. 3. Each antibody is assigned binding sites on M protein. 4. A transfer matrix is defined for each site. It describes the probability for a type of binding to occur at this specific site *i*, provided that the adjacent site *i* + 1 is in a certain state. 5. Through a series of calculations the total binding probability on M protein for any given set of parameters is then determined. See method section for more details on the mathematical framework. **c. Competitive Fab and Fc binding can be predicted with the model using non-competitive experimental values.** As an example of how this model can be applied, an affinity determination is performed on independently measured Fab and Fc binding of human pooled IgG to *S. pyogenes* strain SF370. Data is from three experimental repeats and the error bars indicate standard deviation (*n* = 3, mean *±* SD). The dependent binding is thereafter predicted with the model. **d. Modeling of monoligand Fab- and Fc-specific binding.** By performing calculations with the model for monoclonal antibody fragments with determined binding sites, we show that the model predicts binding as expected. The plots show normalized binding as a function of total IgG concentration. The experimental values were measured using flow cytometry. Data is from three experimental repeats and the error bars indicate standard deviation (*n* = 3, mean *±* SD). SF370 with a GFP plasmid was added to a serial dilution of either IdeS-cleaved *α*M or Xolair and thereafter stained with Fab or Fc specific AlexaFluor647-conjugated F(ab’)2 fragments, respectively. The best fit to the measured binding curves were calculated with the model. Binding affinities extracted from this calculation are given in each of the plots and the ideal binding curve fit serves as a control for the model output. The confidence interval of the affinity estimate is calculated through bootstrapping.

Here, we have developed a biophysical model for describing all of the effects described above. Our implementation of the model is computationally efficient and we provide an easy to use, publicly available Matlab-based software. Computations using our new model shed light on the implications of the IgGFc-binding affinity and demonstrate under which circumstances an effective antibody binding is prominent. We show how our model can be used as a tool for studying the behaviour of various IgG samples by performing in silico simulations in relation to experimentally measured binding values. We use the model here to study the behaviour of specific Fragment antigen-binding (Fab) and Fc binding, pooled IgG and IgG in serum from healthy individual donors. The model is also used to simulate the effect of antibody treatment, by calculating how binding of IgG in serum is altered as specified amounts of monoclonal or pooled IgG is added. Phagocytosis experiments link the altered antibody binding to a physiological function, and demonstrate that it is possible to predict the effect of an IgG treatment with our model.

## Results

### Biophysical modeling of competitive antibody binding to M protein

Here we introduce a a statistical-physics-based theoretical model for the interaction between antibodies and bacterial surface proteins, and show in silico computations with this model in relation to measured binding curves and known affinities. The model describes the binding of IgG to M protein, taking into consideration the binding competition between the Fab and Fc regions as well between different IgG clones. Different IgG binding possibilities on M protein can be described as statistical weights, making it possible to derive the probability of binding to occur at a specific site, and whether it is Fab- or Fc-mediated.

We approached the modeling by implementing the computationally efficient multi-ligand transfer matrix method for competitive binding as described by Nilsson et al and implemented for dual ligand binding to DNA barcodes (21) (25) (26). M and M-like proteins are fairly straight and rigid (18). We therefore model M protein as a one dimensional lattice (**Fig. 1b**). For this linear-protein approximation, M protein is divided into equally large sites *i*. Each antibody is ascribed a size given in a number of sites representing the width it covers on M protein when bound. A polyclonal IgG sample is characterised by its number of clones, a range of affinities to specific sites and the proportion of the bulk concentration of each clone. We can thereby describe a polyclonal IgG sample as a distribution of affinities (16). Each antibody in this distribution is assigned binding sites on the M protein. Additionally, the Fc binding site is defined on the M protein. Every binding possibility has a certain statistical weight that depends on the binding site as well as the affinity and concentration of the IgG clone. The goal is to calculate the probability of a type of binding to occur at a specific site. The probability is given by Equation 1. Equation 1 is numerically evaluated by applying the transfer matrix method. A transfer matrix that includes all states, is defined for each site on M protein. The elements of the transfer matrix describe the statistical weight for a type of binding to occur at a specific site i, provided that the adjacent site *i* + 1 is in a certain state. Through a series of calculations the mean binding probability on M protein for any given set of parameters can then be determined. A more detailed theoretical description is presented in the Methods section and we have made our software publicly available. The computational time required for calculating the binding probability curves such as the ones shown in **Fig. 1** is less than 10 seconds on a standard laptop computer. Altogether we define a system in which a number of antibodies, characterised by their site-dependent affinities, compete for binding to different sites via both Fab and Fc. All possible binding states are identified and the probability of Fab and Fc binding can be calculated using the transfer matrix method.

To test the model, we show experimentally measured independent Fab and Fc binding of pooled human IgG with its corresponding theoretical fit calculated with the model (**Fig. 1c**). The affinities extracted from this model fitting were used to predict the competitive Fab and Fc binding (**Fig. 1c**). Binding of pooled IgG to M protein-expressing bacteria has been measured using flow cytometry. The pooled IgG has been cleaved at the hinge region using the IgG-cleaving enzyme IdeS (28), separating the divalent antibody fragments (F(ab’)_2_) from the Fc fragments prior to being added to the bacterial samples. The measured binding is thus non-competitive between the Fab and Fc region. Independent fits with the model yield affinity estimates for both Fab and Fc binding and binding curves of the competitive binding have been calculated using these. According to this calculation, the Fc binding is out-competed at high total IgG concentrations, which is in line with published findings (22).

By performing calculations with the model for monoclonal antibody fragments with determined binding sites, we show that the model predicts binding as expected. To study Fab and Fc binding independently, IdeS cleaved monoclonal antibodies are used. The Fab binding is studied using a specific antibody to M protein (*α*M, Bahnan et al, manuscript) and a human IgG clone with no specificity to M protein (Xolair) is used for the Fc-binding.

Binding curve fittings, which are performed with the model to values measured by flow cytometry as input data, yield affinity estimates that are verified with an ideal binding curve fitting (**Fig. 1d**). SF370 is added to a serial dilution of either IdeS-cleaved *α*M, or Xolair, and thereafter stained with Fab or Fc specific fluorescently conjugated F(ab’)_2_ fragments, respectively. The determined binding site locations are used to calculate the best fit with the model to the measured binding curve with the *K*_*D*_ value as the unknown variable. The affinity estimates with the model are very similar to those with an ideal binding curve fitting and the calculated Fc affinity is in agreement with previously reported data (30).

### Simulations of Fab and Fc binding capability of M and M-like proteins

To investigate what factors affect the IgG binding of M protein, the effect of the IgGFc affinity on binding is computed at different total IgG concentrations (**Fig. 2a-b**). This was done for two different antibody samples; one with Fab affinities corresponding to that of pooled IgG and one with negligible Fab affinities. According to the first simulation (**Fig. 2a**), the Fc affinity of protein H is precisely at the level to reach maximum Fc binding and minimum Fab binding for the range of physiologically relevant IgG concentrations. However, M protein does not seem to bind Fc at any of these concentrations for the pooled IgG sample. For the antibody sample with no Fab binding (**Fig. 2b**), M protein only binds Fc at serum-like concentrations. This simulation of the IgGFc affinity indicates that protein H has an impact on IgG binding in most host environments, whereas M protein only has an effect in plasma for hosts with very low M protein specificity. This illustrates how calculations with our model can shed light on the implications of IgG binding by bacteria and help determine circumstances under which an effective antibody binding is prominent.

**Fig. 2.**
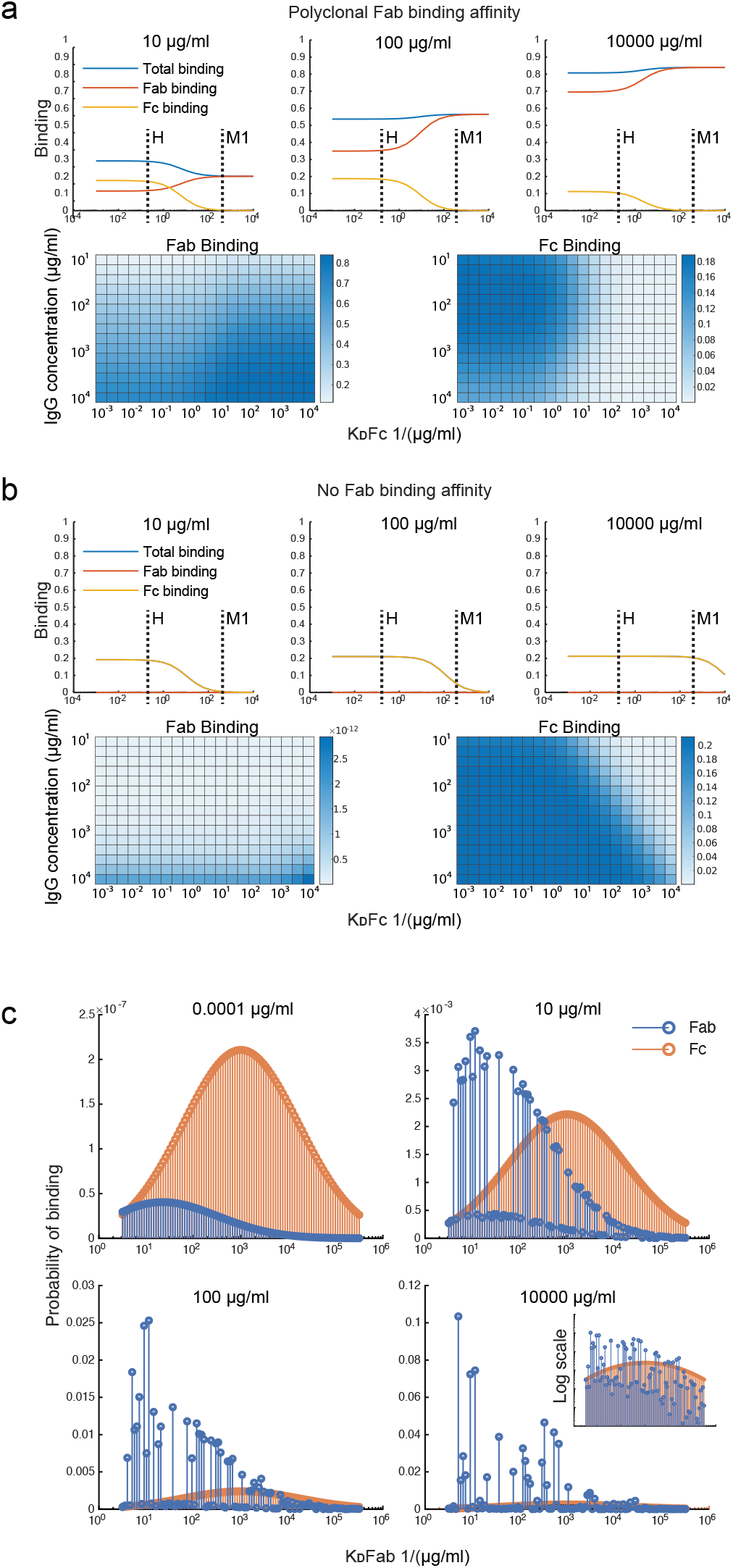
Simulations of Fab and Fc binding capability of M and M-like proteins. **a-b. Simulation of the IgGFc-affinity’s effect on binding.** The effect of the IgGFc affinity on binding has been computed for two different antibody samples; one with Fab affinities corresponding to those of pooled IgG (a) and one with negligible Fab affinities (b). The heatmaps show Fab and Fc binding for different IgGFc affinities and different total IgG concentrations. The plots show binding as a function of Fc affinity at three different concentration points. Fc affinities for protein H and protein M1 are indicated in these plots. **c. Simulation of clonotype-specific antibody binding of a polyclonal IgG sample with normal distributed affinity.** Binding of the different antibody clones is computed for four different total IgG concentrations. The plots show Fab and Fc binding for each of the antibody clones.Fab binding affinities of the different clones are indicated on the x-axis. Note the different ordinate scales.

Simulations were performed to study the clonotype-specific binding of a polyclonal IgG sample (**Fig. 2c**). These calculations indicate that the Fc binding mainly depends on the concentration of the clonotype present whereas Fab binding depends on the clonotype affinity and, at higher concentrations, the epitope location. As some clonotypes may have overlapping epitopes, these antibodies, despite having a higher affinity, may be outcompeted by other clones with slightly higher concentrations. Moreover, according to this simulation a few antibody clones constitute a large proportion of the total Fab binding at plasma-like concentrations. The clonotype-specific simulations illustrate that epitope location and competition between clones are important for the M protein-IgG interaction.

### Binding of polyclonal IgG samples

We wanted to apply our model to situations that are relevant for both mechanistic understanding as well as for potential treatment scenarios, meaning we had to expand it to polyclonal IgG. Simulations are presented here for pooled human IgG (IVIgG) and IgG in serum from healthy donors. The effect of adding specified amounts of specific or pooled IgG to human serum is computed with the model. IVIgG is a complex polyclonal IgG sample that presumably contains a large number of antibody clones with a broad spectrum of affinities to M protein. Fab binding of IVIgG to M protein is estimated by measuring binding curves to SF370 wild type as well as an M protein mutant (**Fig. 3a**). The difference between these binding curves should correspond to Fab-binding of IVIgG to M protein. The model fitting yields a mean affinity of 10^2.2*±*0.1^μg/ml and a range in log_10_ of 0.55 0.8. Figure **Fig. 3a** shows a heatmap over the error of the fit for the mean and range of the affinity distribution, illustrating the sensitivity of the model to these parameters. We have thus measured binding of IVIgG to M protein and shown how we can estimate affinity values of this polyclonal sample using the model.

**Fig. 3.**
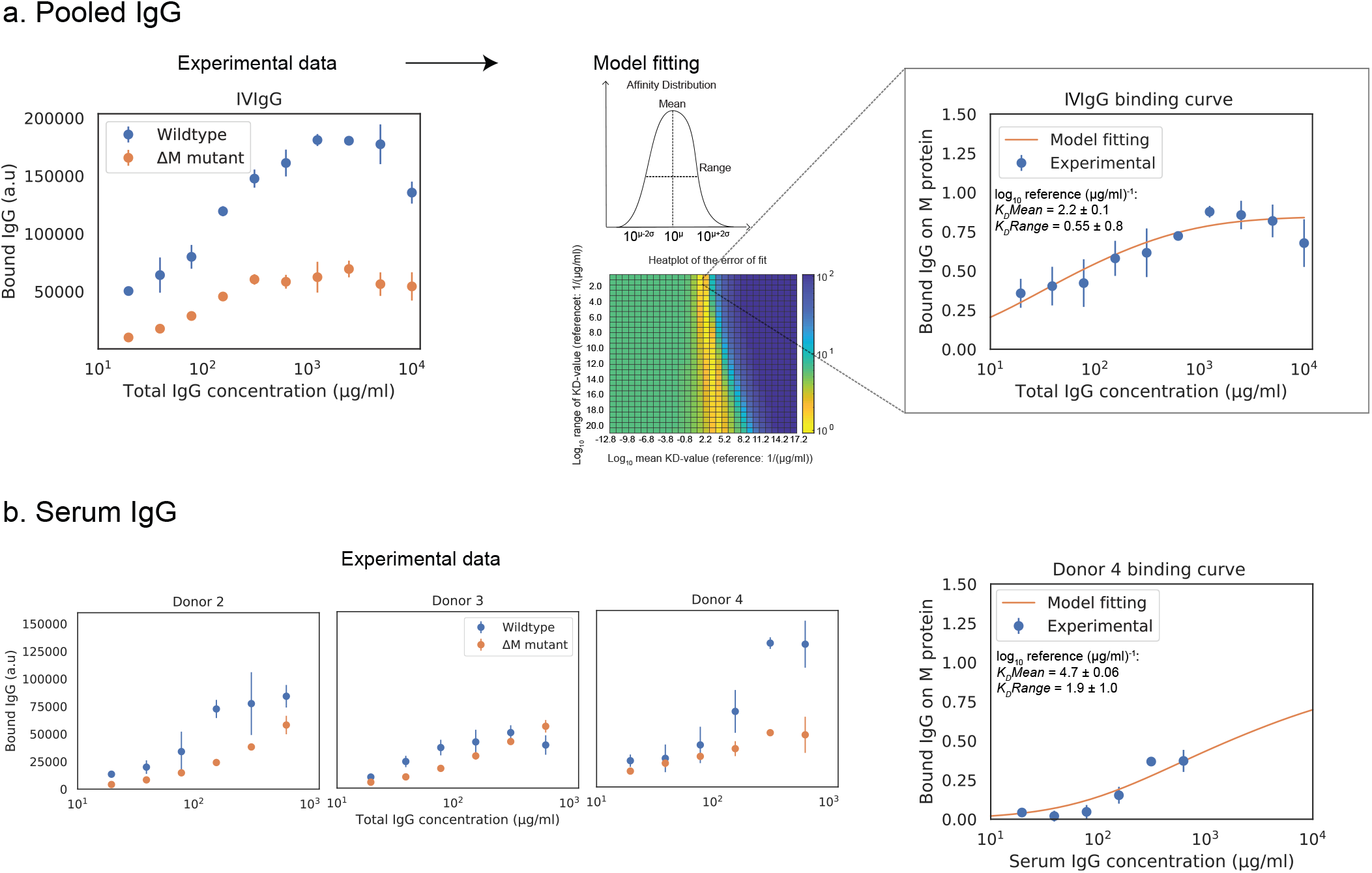
Binding of polyclonal IgG samples. Simulations of polyclonal IgG samples are presented here for pooled human IgG and IgG in serum from healthy donors. **a. The behaviour of pooled human IgG is studied with the model.** Binding curve of IVIgG to SF370 wild type and M protein mutant The Fab binding of IVIgG to M protein is estimated to be the difference between binding of IVIgG to SF370 wild type and the M protein mutant. The data shown is from three experimental repeats and the error bars indicate standard deviation (*n* = 3, mean *±* SD). Binding of IVIgG to M protein with model fitting Assuming a broad distribution of affinities present in IVIgG, we calculate the mean and range of this distribution for the experimentally measured binding of IVIgG to M protein. The plot shows normalised binding as a function of total IgG concentration for the measured binding of IVIgG to M protein together with the calculated best fit to the model output. Data is from nine experimental repeats and the error bars indicate standard deviation (*n* = 9, mean *±* SD). The heatmap shows the error of the fit for the mean and range of the affinity distribution for a broad range of values. **b. Binding curves of donor serum to SF370.** The behaviour of IgG in human serum from healthy donors is studied with the model. As with IVIgG, the Fab binding of serum IgG to M protein is estimated to be the difference between binding of serum IgG to SF370 wild type and the M protein mutant. A model fitting is performed for one of the serum binding curves. The bottom plot shows normalised binding as a function of total IgG concentration for the measured binding of serum IgG to M protein together with the calculated best fit to the model output.

### Model fitting and titration simulations of monoclonal and pooled IgG

To test this model in host-environment like conditions, binding of IgG in serum from individual donors was studied. As with IVIgG, Fab binding of IgG to M protein in serum from healthy individual donors is measured and simulated with the model (**Fig. 3b**). Interestingly, the Fab binding curves of the serum samples show large variations in binding to M protein. A model fitting performed for one of the serum binding curves is shown in **Fig. 3b** together with its affinity estimates. We next calculate how this binding curve is altered when specified amounts of *α*M or IVIgG is added to the serum, corresponding to different routes of administration (**Fig. 4**). We show simulations of how the binding curve for donor 4 is altered as the present IgG is increased by a range of percentages of either IVIgG or *α*M (**Fig. 4**). A proportional increase in present IgG illustrates the case of intravenous IgG administration with an equally proportional dispersion to tissue environments. For the clinical use of IVIgG as treatment for invasive infections, about 200% of serum IgG (~ 10 mg/ml) is typically administered (13). Additionally, we show simulations of how this binding curve is altered as specific concentrations of antibody are added at each IgG concentration. An increase of present IgG by a specified concentration (**Fig. 4**) can represent administration by injection to an infected area or a non-proportional delivered local dose. These simulations illustrate how we can use this model to represent various pathophysiological scenarios of delivered local IgG dose.

**Fig. 4.**
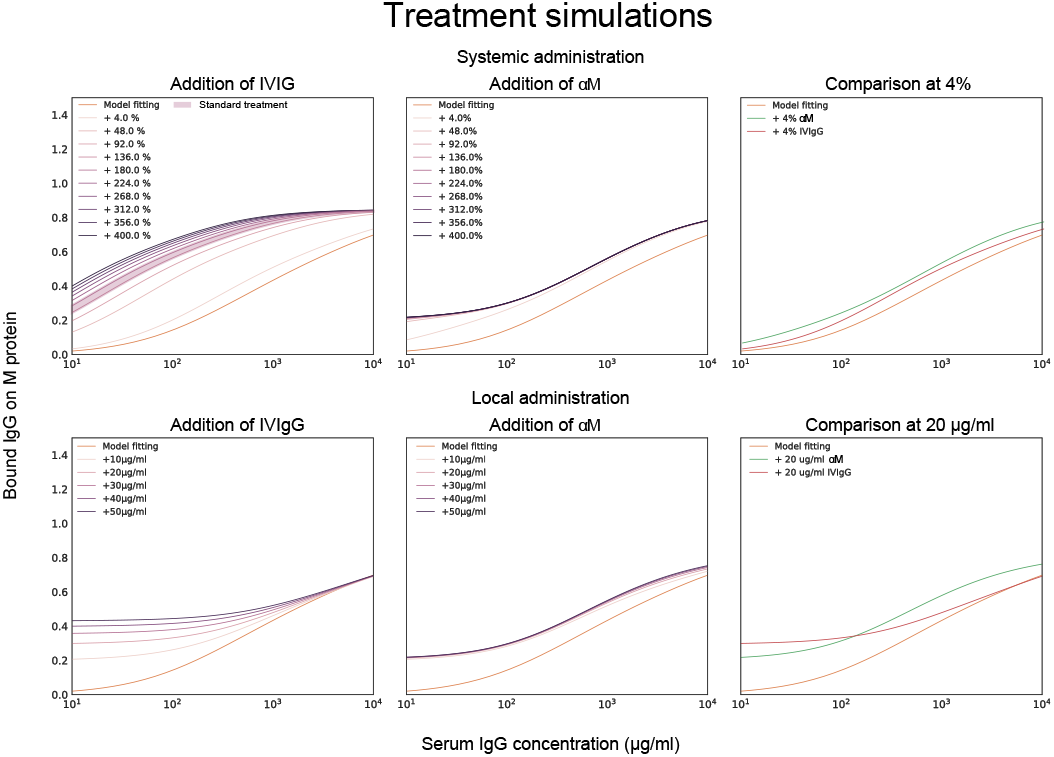
Model fitting together with titration simulations of monoclonal and pooled IgG. It is calculated how the binding curve for donor 4 is altered when specified amounts of *α*M or IVIgG is added to the serum. Previously calculated affinities for *α*M and IVIgG are used for this simulation. The top three simulated plots show how the binding curve for donor 4 is altered as the IgG present is increased by a range of percentages of either IVIgG or *α*M. The bottom three plots show how the binding curve shifts as a specific concentration of antibody is added at each given serum IgG concentration. The case of 4% and 20 μg/ml increase are shown for *α*M and IVIgG together for comparison in the two rightmost plots.

### Phagocytosis in serum with monoclonal or pooled IgG

To evaluate how added monoclonal or pooled IgG affects phagocytosis of *S. pyogenes* in serum, 20 μg/ml of either *α*M or IVIgG is added to 5% of serum (**Fig. 5**). This corresponds to a 4% increase of antibody in full serum. According to the simulations, at 5% serum 20 μg/ml of *α*M gives higher binding than the same amount of IVIgG (**Fig. 4**). The phagocytosis experiments are performed according to a standardised assay based on normalisation of persistent association of prey and phagocyte (6). Association measured with Xolair-coated bacteria serves as a baseline of non-Fc-mediated association. An increase in association is seen as *α*M or IVIgG is added to the serum. The association is slightly higher for *α*M than for IVIgG (MOP_50_= 29.65 7.6 (*mean SD*) for *α*M and MOP_50_= 39.75 ± 11.8 for IVIgG). This difference is however not statistically significant (*p* = 0.3). Additionally, the proportion of phagocytes that internalise seem to be increased at higher MOP when *α*M or IVIgG is added. Considering the area under curve (AUC) as a cumulative measure of internalisation, we see that serum with *α*M has the highest total internalisation, and that adding IVIgG to serum does not considerably increase total internalisation. We expect an increase in Fab binding to yield more Fc-mediated association of cells and bacteria. Overall, we find that the presented increase in association and proportional internalisation agrees well with the predicted results from the model.

**Fig. 5.**
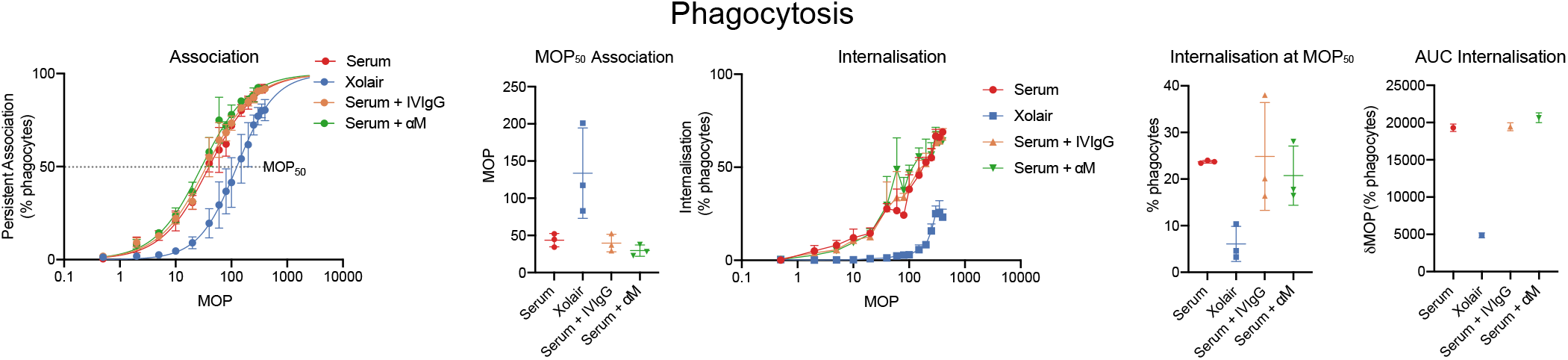
Phagocytosis experiments. Association of phagocytes and bacteria treated with different antibody samples are shown to the left. These curves depict the percentage of present phagocytes that have associated with at least one bacteria as a function of the multiplicity of prey (MOP). The data shown is from three experimental repeats and the error bars indicate standard deviation (*n* = 3, mean *±* SD). The MOP at which 50% of phagocytes have associated (the MOP_50_) is illustrated in a bar plot for the different antibody treatments. The internalisation curves depict the percentage of phagocytes that have internalised one or more bacteria as a function of MOP and the internalisation rates at MOP_50_ are shown in a separate plot. The rightmost plot depicts the area under curve (AUC) of the internalisation curves.

## Discussion

In this article we have presented a framework for a competitive antibody binding model together with its potential applications. Our implementation of the model is computationally efficient and we provide an easy to use, publicly available Matlab-based software. We have experimentally verified the model and have shown how this model can be used to determine binding characteristics of both monoclonal and polyclonal IgG samples as well as IgG in a serum environment. Additionally, the model has been used to perform in silico computations to understand the inherent behaviour of the IgG – M protein interaction, such as the effect of the IgGFc affinity on the overall binding. Ultimately, we have illustrated how this model can be used as a tool to predict the effect of antibody treatments by simulating how binding of IgG in serum is altered as specified amounts of monoclonal or pooled IgG is added.

The model that we have developed takes into account aspects of the protein-antibody interaction that hitherto have not been considered. The concentration dependency of the IgGFc-binding is in line with previous results (22). The model behaves as an ideal binding curve for monoligand binding. Finally, we have shown that functional assays are consistent with simulations of antibody treatment. Due to the fact that the competitive binding model incorporates parameter values that could not be determined experimentally, simulations with the model require certain assumptions regarding these. Specifically, the number of clonotypes in pooled IgG or serum and how their affinities are distributed is generally unknown and has therefore been based on measurements of specific antibodies (17) (16). While the model agrees well for these conditions, the values extracted from the model need to be understood in the context of the assumptions made regarding these parameters.

The competitive binding model could be improved by incorporating other relevant ligands of M protein that are present in the host environment. The framework is designed as such that it can be extended to include other human proteins that bind M protein, such as fibronectin, fibrinogen, C4BP and factor H. Another possible extension of this model would be to calculate IgG binding to other bacterial surface proteins. Protein G of group C and G streptococci and protein A of *S. aureus* are globular proteins with five and three IgGFc-binding regions respectively. A more accurate model for binding on the surface of globular proteins could be attained through two-dimensional transfer matrix calculations.

Prediction of the competitive binding of pooled IgG gives insight into the previously observed concentration dependency of IgGFc-binding (22). The Fc-binding affinity is constant and there exists a spectrum of Fab-binding affinities to multiple epitopes on M protein. Thus, with an increase of concentration of antibodies, the probability of high-affinity Fab-binding increases to an extent where the Fc-binding is outcompeted. This together with the previous findings indicates that IgGFc-binding by bacteria has evolved to execute its function most efficiently in IgG-poor environments such as saliva, and may explain why severe invasive infections in blood are so rare.

Simulations illustrate how we can use this model to represent various pathophysiological scenarios of delivered local IgG dose and give an insight into what these scenarios might implicate. According to these calculations, whether pooled IgG or a specific mAb gives rise to the highest Fab binding depends on both the amount and route of administration. Specifically, 4% antibody treatment with mAb gives more binding than IVIgG. However, the inverse is true at 48% administration and above. In fact, the binding increase of the mAb treatment saturates at relatively low percentages while the binding of IVIgG continues to increase at higher concentrations. A probable explanation for this is that there exists only one epitope for mAb whereas IVIgG has several clones with different epitopes (22), thus the mean binding on M protein can increase to a higher extent with IVIgG than with mAb. A specified local dose as treatment gives rise to higher binding with IVIgG than with mAb in low IgG environments. At high concentrations treatment with a specified dose of mAb increases binding more than with IVIgG. Even though the mean affinity of IVIgG is lower than that of mAb, as per our calculation, there exist antibody clones in IVIgG with higher affinity than mAb. These results suggest that high affinity IgG clones have the most effect in low IgG environments. The increase in Fab binding with the treatments should increase Fc-mediated association of cells and bacteria.

The results from the phagocytosis experiments are consistent with the simulation. Internalisation of bacteria by cells link the antibody binding to a physiological effect. Altogether these results indicate that there are several factors to consider when selecting an antibody treatment. Even though pooled IgG yields higher binding in certain scenarios, it is more difficult to characterise the effect of this binding than with mAbs. Furthermore, the delivered local dose to infected regions needs to be considered. In the future, this model could potentially be used to determine the most efficient antibody treatment and route of administration for a given infection, as well as assist in identifying the therapeutic range of an antibody treatment for a specific patient.

## Method

### Competitive antibody binding model

Protein-protein binding is in essence an interaction resulting in a stable complex with lower free energy than when the proteins are unbound. The binding is formed through cumulative attractive forces between the atoms in the proteins and the complexity of these large molecular energy systems requires them to be described statistically.

By treating different binding possibilities on M protein as statistical weights and using statistical physics, a formula for the probability of a type of binding to occur at a specific site could be derived. The straight and rigid structure of M protein enables us to model it as a one-dimensional lattice. This lattice is divided into *N* number of sites, each denoted *i*. Consider *S* number of antibody clones denoted *s*(*s* = 1,2,….*S*) competing for binding to the sites on M protein. The antibodies can bind via *L*(*l* = 1,2…*L*) different regions to M protein. For antibodies *L* is set to 3 wherein *l* = 1,2 correspond to binding via Fab and *l* = 3 binding via Fc. The type of binding *l* for antibody clone *S* is characterised by a site-dependent binding constant *K*_*s,l*_(*i*) that relates to its binding affinity. *λ* is the number of sites *i* an antibody covers when bound to a specific site on the lattice ((**Fig. S1**). Each possible binding state has a statistical weight which is given by the binding constant as well as the present free concentration of the antibody clone *c*_*s*_, as *c*_*s*_*K*_*s,l*_(*i*) (**Fig. S1**). We aim to calculate the probability *p_s,l_*(*i*) of a type of antibody binding to occur at a specific site on M protein. The probability *p*_*s,l*_(*i*) is

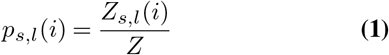

where *Z* is the partition function and *Z*_*s,l*_(*i*) is the sum over all allowed Boltzmann weighted states for site i.

To numerically evaluate 1 we implemented the multi-ligand transfer matrix method for competitive binding as described by Nilsson et al (21). The number of configurations that need to be considered in order to evaluate 1 increases exponentially with *N*. Using the transfer matrix method the probability resulting from all these configurations can be calculated efficiently. The transfer matrix method is applied to calculate *Z* and *Z*(*i*) for this system. There exists *M* = 1 + *lλ*S* number of possible states for each site *i*. An *M* × *M* transfer matrix *T* (*i*) is defined for each site *i*. *T* (*i*; *m, m*′) gives the statistical weight of *i* being in state *m* provided *i* + 1 is in state *m*′. A specific order that is chosen for the possible states together with their respective statistical weights is illustrated in **Fig. S2**. The centermost region of the antibody is assigned the statistical weight for binding. Explicitly, the elements of the transfer matrices can be described as follows:

- Site *i* and *i* + 1 are both empty: *T* (*i*; 1, 1) = 1
- Site *i* is empty while *i* + 1 is covered by the leftmost segment of antibody *s*: *T* (*i*; 1, 1 + *λs*) = 1, If *i* + 1 is covered by the leftmost segment of any antibody, it is only possible for *i* to be either empty or covered by the rightmost segment of any other antibody.
- Site *i* is covered by the rightmost segment of antibody *s* and site *i* + 1 is empty *T* (*i*; 2 + *λ*(*s* − 1), 1) = 1
- Both site *i* and *i* + 1 are covered by antibody *s*; excluding all cases in which the centermost region of *s* is bound to site *i*: *T* (*i, m* + 1*, m*) = 1.
- Site *i* is covered by the rightmost segment of antibody *s* and site *i* + 1 is covered by the leftmost segment of another antibody *s′*′: *T* (*i*; 2 + *λs* − 1, 1 + *λs′*′) = 1
- Site *i* is bound to the centermost region of antibody *s*, *i*+1 is hence covered by the following part of antibody *s*. *T* (*i*; 1 + *λs, λs*) = *c*_*s*_*K*_*s,l*_(*i*)

The cases described above cover all possible states and therefore the remaining elements in the transfer matrix equal 0. To illustrate a simple case for the following part of this section, we show the transfer matrix for one type of antibody *S* = 1, *l* = 3 and *λ* = 3. *M* is thus 10 and each transfer matrix takes the form

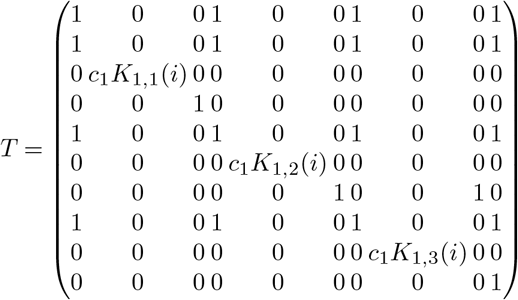

*Z*_*s,l*_(*i*) and thereby *p*_*s,l*_(*i*) can be calculated using the transfer matrices. The partition function *Z* is

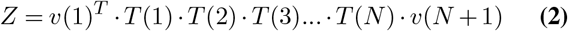

where *v*(1) and *v*(*N* + 1) are column vectors of length *M*, and consist of all allowed states of the first and last site on M protein, considering the geometry of the antibody and bacterial surface. For our example *v*(1) would be

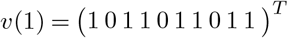

and assuming the last site is fixed to the bacterial surface *v*(*N* + 1) equals

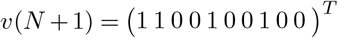

thus disallowing binding that would be physically blocked by the bacterial surface. The sum over all allowed Boltzmann weighted states *Z*_*s,l*_(*i*) is described as

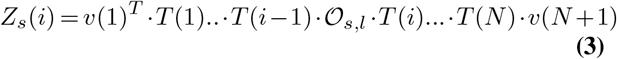

where the projection operator 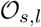 has been introduced. 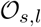 selects states wherein a particular type of binding occurs. For instance, the projection operator for all states associated with Fab binding, i.e. states for which *l* = 1,2 and *s* = 1,2..*S*, 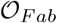 is, as per the previously defined choice of enumeration

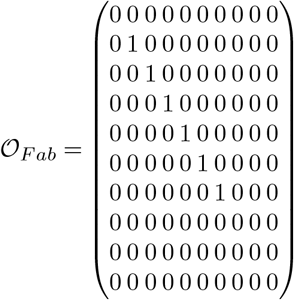

In the same manner 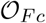 is 0 for all elements except for those corresponding to Fc-binding, i.e. the states for which *l* = 3 and *s* = 1,2..*S*. The probabilities *p*_*Fab*_(*i*) and *p*_*Fc*_(*i*) can thus be calculated for all sites on the M protein. The mean probability that sites are occupied by Fab or Fc bound antibodies for the bacterial protein as a whole is then

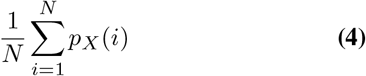

where X = Fab or Fc.

### Implementation of model

In the following section we describe how the model has been implemented for the simulations in this article and present the reasoning behind the adopted values for parameters that could not be determined experimentally. The parameters for this model are *N*, *λ*, *K*_*s,l*_(*i*), *c*_*s*_ and *S*.

To start with, for all simulations of polyclonal IgG each antibody *s* is assigned one binding site with an arbitrary location. Since the binding site locations of specific antibodies in polyclonal samples are generally unknown, these have been generated through randomisation. Antibodies typically recognize a particular sequence of amino acids and thus have high affinity to a specific epitope. There may nevertheless occur binding to other epitopes on the bacterial protein with lower affinities. This aspect is incorporated in the formalism of the model. However we have made the simplification in the implementation that each antibody *s* only binds to a specific site *i*.

As the ratio between *N* and *λ* corresponds to the proportion of M protein that an antibody covers when bound, these parameters have been based on the width of the antibody relative to the length of the bacterial protein. *λ* was thus set to 7 and *N* set to 33 (22) and increasing *N* and *λ* proportionally from this did not affect the result **Fig. S4**.

A polyclonal IgG sample is defined here by its number of clones *S*, a range of affinities to specific sites *K*_*s,l*_(*i*) and the concentration of each clone *c*_*s*_. Polyclonal samples such as pooled IgG presumably contain a large number of antibody clones with a broad spectrum of affinities to M protein. Previous studies suggest that serum from an immunised individual can have >100 specific IgG clones (17). Additionally, due to the arbitrary assignment of binding sites, computations with the model are more stable for an increasing amount of IgG clones **Fig. S3**. Therefore all calculations have been performed for 50-200 IgG clones. Affinity measurements of 1,450 monoclonal antibodies show that the exponentials of the *K*_*D*_ values are normally distributed, suggesting that the dissociation constant for IgG is an inherently exponential measurement (16). We have therefore characterised antibody affinities in polyclonal samples as a normal distribution of the *K*_*D*_ exponents. The range and mean affinities of these distributions are determined by fitting to experimentally measured values. All fittings to measured data have been done using a weighted least squares method. The free concentration of antibodies is attained through an iterative minimisation with the total concentration as initial value. The implementation of the model has been written in MATLAB, and is available through GitHub (https://github.com/nordenfeltLab/ComputeTM).

### Experimental methods

#### Bacterial culturing conditions

*S. pyogenes* strain SF370 wildtype and ΔM mutant (1) were cultured overnight in THY medium (Todd Hewitt Broth; Bacto; BD, complemented with 0.2% (w/v) yeast) at 37 °C in an atmosphere supplemented with 5% CO_2_. Strain SF370 expresses M1 protein on its surface and is available through the American Type Culture Collection (ATCC 700294)(8). For the binding curve measurements SF370 expressing GFP was cultured with 1 μg/ml erythromycin. The bacteria were harvested at early log phase and washed twice with PBS. For the assays in which live bacteria were used 0.2% glucose was added to the PBS.

#### Opsonisation of bacteria

Human serum collected from 4 healthy individuals was heat-inactivated at 56 °C for 30 min and stored in aliquotes at −20 °C. Intravenous immunoglobulin (IVIgG; Octagam, Octapharma) is IgG pooled from >3500 healthy donors. Xolair (Omalizumab, Novartis) is a humanised monoclonal IgG that is IgE-specific, and thus only binds to M protein via Fc. *α*M is an M protein specific antibody (Bahnan et al manuscript in preparation). Based on reference values, the concentration of IgG in full serum was set to 10 mg/ml (27).

Serum and all antibody samples were incubated with IdeS (1 μg/ml to 5 μg/ml depending on the maximum IgG concentration of the serial dilution) at 37 °C overnight. Bacteria were sonicated (VialTweeter; Hielscher) for 0.5 minutes to separate any aggregates and incubated with serial dilutions of serum, IVIgG, Xolair and *α*M for 30 minutes on shake (400 rpm) at 4 °C to minimise metabolic activity of the bacteria. Unbound antibodies were washed away after incubation with PBS (3220g 3 min) 2-4 times, depending on the maximum IgG concentration of the serial dilution.

### Measuring binding curves

#### Antibody staining

Bacterial samples opsonised with antibodies were stained with either fluorescently labelled IgGFab or IgGFc specific F (ab’)_2_ fragments (Alexa Fluor 647-conjugated F(ab’)_2_ Fragment Goat Anti-Human IgGFab and Alexa Fluor 647-conjugated F(ab’)_2_ Fragment Goat Anti-Human IgGFc).

#### Flow cytometry acquisition

Flow cytometry acquisition was performed with CytoFLEX (Beckman-Coulter) with 488- and 638 nm lasers, and 525/40 FITC and 660/10 APC filters. The bacteria were gated by SSC-H and the GFP signal using control sample with only bacteria. The antibody signal background gating was performed using control with only bacteria and the antibody staining (Alexa Fluor 647-conjugated F(ab’)_2_ Fragment Goat Anti-Human IgGFab or IgGFc). Acqisition was set to run until at least 20 000 events were collected in the target population or 200 μl of sample had run through.

#### Analysis

The data was analysed using MATLAB and files are available on GitHub. For serum and IVIgG samples, binding to M protein was derived as the difference between mean wildtype and ΔM binding. The standard deviation is the square root of the sum of the variances of wildtype and ΔM. Binding curve values were normalised to an interpolated saturation level prior to being evaluated with the model implementation. Measured binding curves are shown as mean and standard deviation of data points as described in the figure legends above. Affinity values were derived by minimising the mean squared error of the model output and measured data using a MATLAB minimisation function. *K*_*D*_ values extracted from an ideal binding curve were derived in the same manner for a monoligand dose-response curve together with the measured data. The accuracy of predicted affinity estimates was calculated using the bootstrap method (7) and is the confidence interval calculated from 50 resamplings of the measured data.

### Phagocytosis assay

#### Microbe strains

*S. pyogenes* strain SF370 was cultured at 37 °C with 5% CO_2_ atmosphere in 50 ml THY-medium (Todd Hewitt Broth; Bacto; BD, complemented with 0.2% (w/v) yeast). The bacteria were harvested at optical density OD600 0.5.

#### Cell lines

THP-1 cells (ATCC TIB-202, male) were maintained in Roswell Park Memorial Institute medium 1640 (Sigma-Aldrich) supplemented with 10% fetal bovine serum (Gibco) and 2 mM GlutaMAX (Life Technologies). Cells were grown at 37 °C in a 5% CO_2_ atmosphere. The cell density was kept between (0.2 1.0) × 10^6^ cells/ml with a viability over 95% and split to 0.5*x*10^6^ cells/ml the day prior to experiment.

#### Heat-killing of bacteria

After harvesting the bacteria, they were washed 2 times with 10 ml PBS (Gibco) at 4000 x g for 10 min and resuspended in 1 ml PBS. Heat-killing was done at 80 °C for 5 min on a vigorously shaking heat block, followed by cooling the bacteria rapidly on ice.

#### Labeling of bacteria

Heat-killed bacteria were stained with 5 μM Oregon Green 488-X succinimidyl ester (Invitrogen) at 37 °C with gentle rotation and protected from light for 30 min. This was followed by a change of buffer to sodium carbonate (0.1 M, pH 9.0) and staining with the pH-sensitive dye CypHer5E (General Electric) 20 μg/ml for 2 h in room temperature under gentle rotation in the dark. The samples were washed two times with Na-medium (5 min, 8000 x g, swing-out rotor). The stock of labeled bacteria was stored up to 5 days in fridge protected from light and was used for all phagocytosis experiment. Prior to opsonization bacteria were sonicated until (VialTweeter; Hielscher) any large aggregates of bacteria were disperse, which was confirmed by microscopy. Staining was checked by flow cytometry (CytoFLEX, Beckman-Coulter), for CypHer5E, 1 μl of sodium acetate (3 M, pH 5.0) was added to confirm pH-sensitivity.

#### Opsonization

The concentration of labeled bacteria was measured by flow cytometry. During opsonization the concentration was kept at 400000 bacteria/μl. The opsonization was performed with 5% patient heat-inactivated sera in Na-medium (5.6 mM glucose, 127 mM NaCl, 10.8 mM KCl, 2.4 mM KH2PO4, 1.6 mM MgSO4, 10 mM HEPES, 1.8 mM CaCl_2_; pH adjusted to 7.3 with NaOH). For two samples the sera were supplemented with either intravenous immunoglobulin (IVIG, Octagam, Octapharma, 20 μg/ml) or *α*M (Bahnan et al manuscript in preparation, 20 μg/ml). As negative control the non-specific antibody solution Xolair (Novartis, 500 μg/ml) was added to Na-medium without sera. Opsonization occurred at 37 °C with gentle shaking and protected from light for 30 min.

#### Phagocytosis assay

The cell density was kept at 0.5 × 10^6^ cells/ml on the day of experiment. Medium was changed to Na-medium (10 min, 4696 x g swing out rotor). The concentration of THP-1 cells was measured prior to phagocytosis by flow cytometry and adjusted to 2000 cells/μl, amounting to 100 000 cells per well. A 96-well plate was prepared with the bacteria with a multiplicity of prey (MOP) of 0-400. 50 μl of THP-1 cells were thereafter added on ice, resulting in a final volume of 150 μl. The plate was directly transferred to 37 °C on a heating-block with moderate shaking protected from light or kept on ice as control for internalization. Phagocytosis was haltered by putting the samples on ice for at least 15 min before data acquisition. Three independent experiments were performed.

#### Data acquisition

Flow cytometric acquisition was performed using CytoFLEX (Beckman-Coulter) with 488 nm and 638 nm lasers and filters 525/40 FITC and 660/10 APC. Threshold was set at FSC-H 70 000 for phagocytosis and for bacteria FSC-H 2000 and SSC-H 2000. Gain was kept at 3 for FITC and 265 for APC. Acquisition was set to analyze at least 5 000 events of the target population with a velocity of 30 μl/ min taking approximately 60 min for all samples. Throughout the data acquisition the plate was kept on an icecold insert.

#### Data analysis

Flow cytometry data were analyzed using FlowJo version 10.2 (Tree Star). The THP-1 cells were gated on forward (FSC) and side scatter (SSC) height, then doublets were excluded by gating on FSC-H versus FSC-A. THP-1 cells positive for Oregon Green-signal were defined as associating cells, and of those THP-1 cells, cells positive for CypHer5E-signal were defined as internalizing cells. Associating and internalizing cells are expressed as the percentage of the whole phagocytic population. The data was further analyzed using the PAN-method (6).

## Author contributions

VKA developed the antibody binding model, designed and performed experiments, analyzed data, and wrote the manuscript. TdN designed phagocytosis experiments and analyzed data. MS performed phagocytosis experiments and analyzed data. TA supervised the development of the antibody binding model. PN supervised the project and wrote the manuscript. All authors reviewed the final manuscript.

## ACKNOWLEDGEMENTS

VKA and TdN was funded by the Royal Physiographic Society. PN was funded by the Swedish Research Council, the Crafoord Foundation, and the Knut and Alice Wallenberg Foundation.

---

**Fig. S1.**
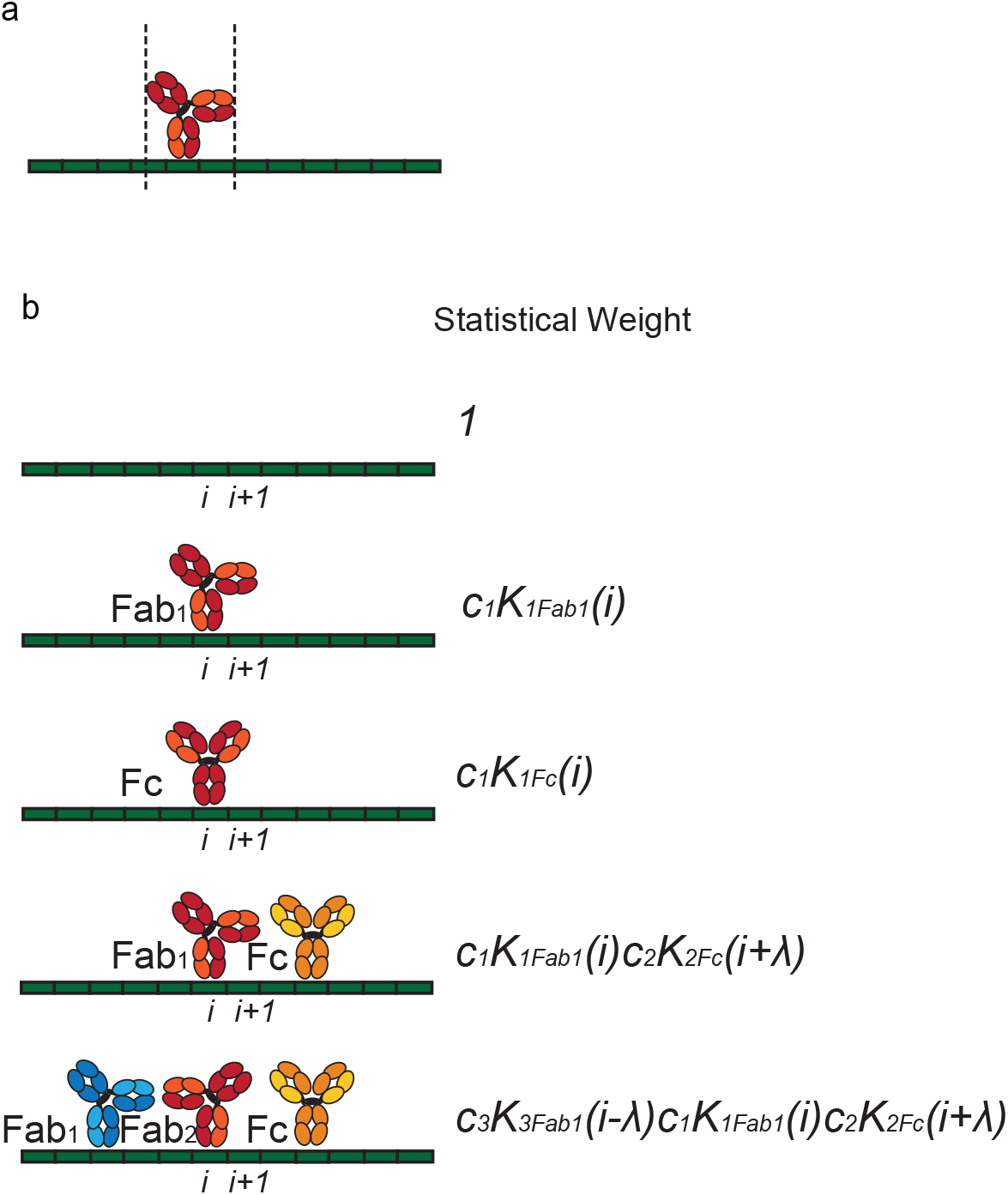
a. A schematic of antibody *s* bound to a site on the linear bacterial protein while covering two additional sites, thus illustrating the case *λ* = 3 b. Examples of statistical weights. Different clonotypes are shown in different colours. The state in which no antibodies are bound to the bacterial protein has the statistical weight 1, whereas the statistical weight for a bound antibody depends on its present concentration *c*_*s*_ and the site and fragment specific binding constant *K*_*F ab/F c*_(*i*)

**Fig. S2.**
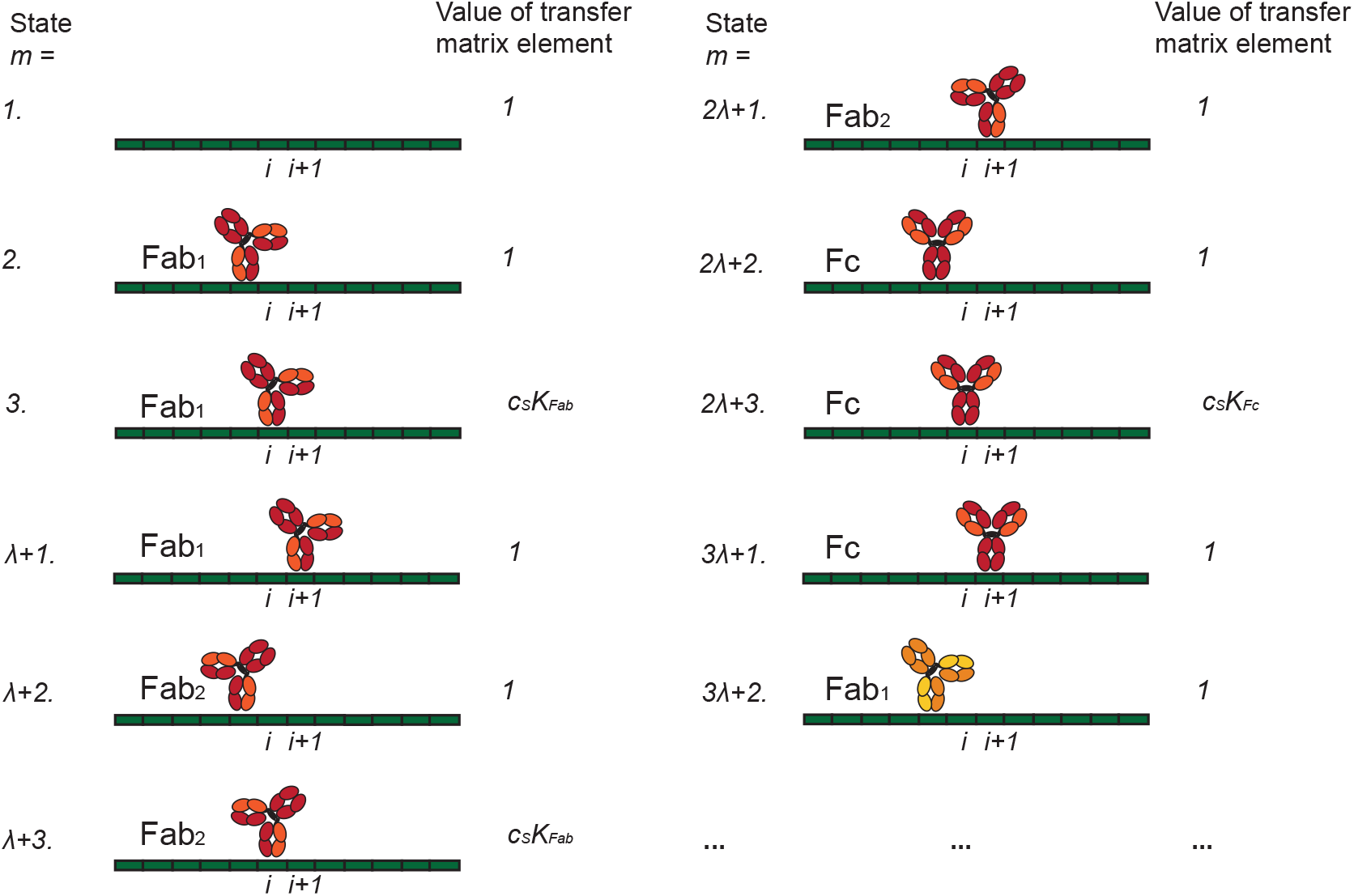
The choice of enumeration adopted for our transfer matrices. The binding states *m* are illustrated together with their transfer matrix values for site *i*. All states for the first antibody clone *s* = 1 (in red) are shown, as well as the first state for the antibody *s* = 2 (in yellow). The states of the subsequent antibody clones follow the same pattern.

**Fig. S3.**
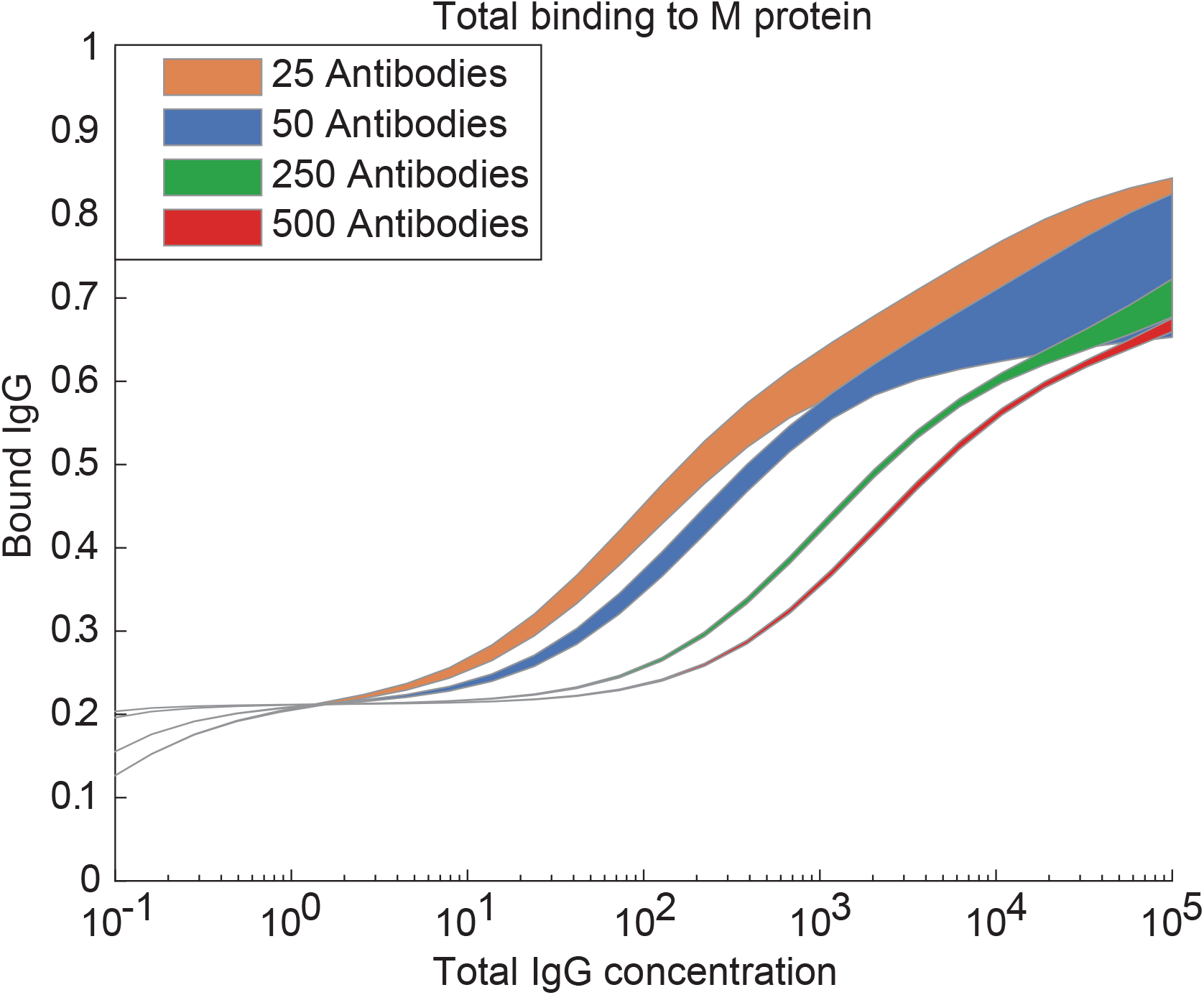
The standard deviation calculated from 10 computed curves of total binding to M protein, each with a different set of randomly generated epitopes, for polyclonal IgG samples with different numbers antibody clones *S*. The simulation indicates that computations with the model are more stable for an increasing *S*.

**Fig. S4.**
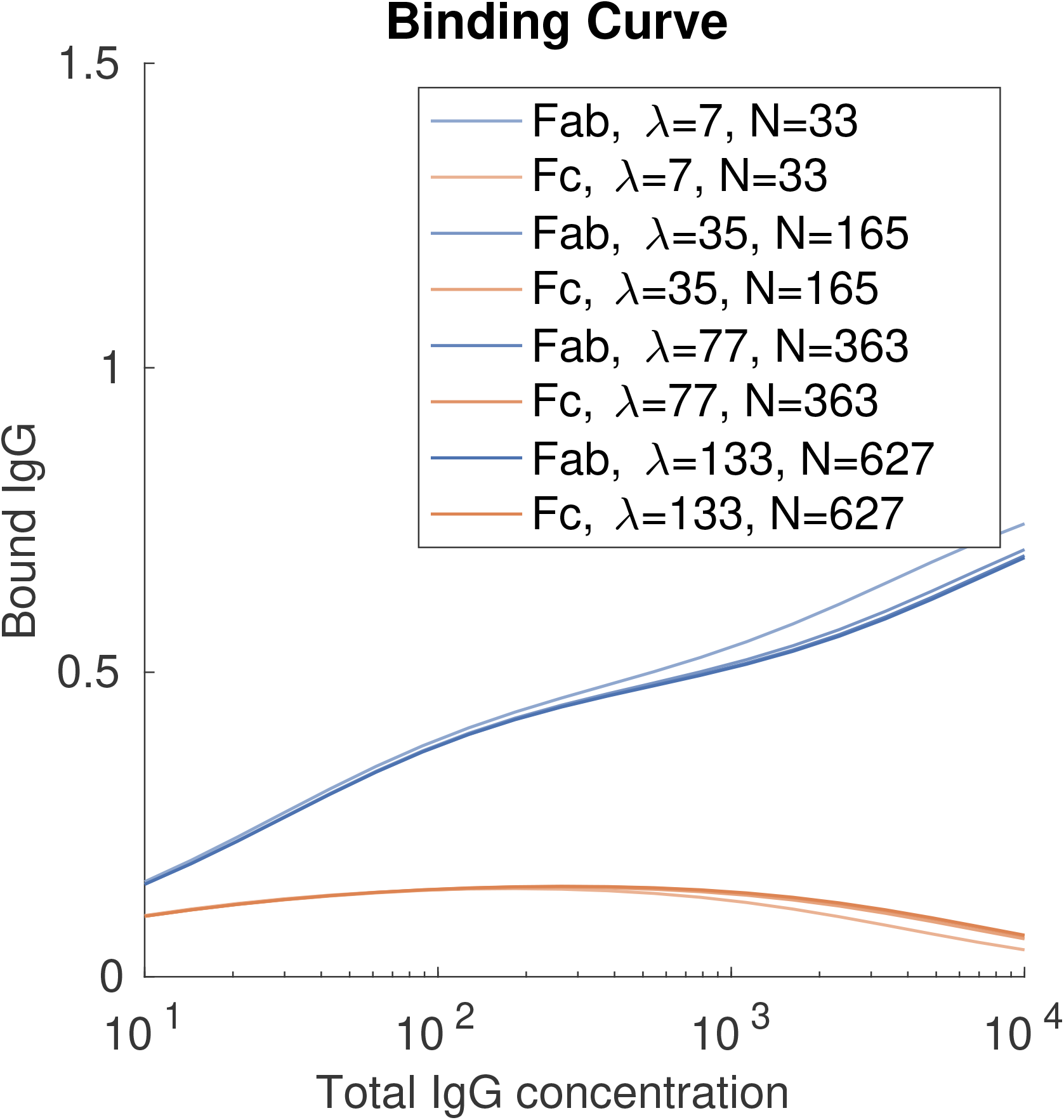
Fab and Fc binding curves calculated for a polyclonal IgG sample with values set as for pooled IgG. *N* and *λ* are increased proportionally from the set values 33 and 7 respectively, as described in the methods section.

## Bibliography

1. E. L. Abbot, W. D. Smith, G. P. S. Siou, C. Chiriboga, R. J. Smith, J. A. Wilson, B. H. Hirst, and M. A. Kehoe. Pili mediate specific adhesion of streptococcus pyogenes to human tonsil and skin. Cell Microbiol, 9(7):1822–1833, jul 2007.

2. L. Björck and G. Kronvall. Purification and some properties of streptococcal protein g, a novel IgG-binding reagent. J Immunol, 133(2):969–974, aug 1984.

3. J. R. Carapetis, P. Jacoby, K. Carville, S.-J. J. Ang, N. Curtis, and R. Andrews. Effectiveness of clindamycin and intravenous immunoglobulin, and risk of disease in contacts, in invasive group a streptococcal infections. Clin Infect Dis, 59(3):358–365, aug 2014.

4. J. R. Carapetis, A. C. Steer, E. K. Mulholland, and M. Weber. The global burden of group a streptococcal diseases. Lancet Infect Dis, 5(11):685–694, nov 2005.

5. J. Darenberg, N. Ihendyane, J. Sjölin, E. Aufwerber, S. Haidl, P. Follin, J. Andersson, A. Norrby-Teglund, and S. S. Group. Intravenous immunoglobulin g therapy in streptococcal toxic shock syndrome: a european randomized, double-blind, placebo-controlled trial. Clin Infect Dis, 37(3):333–340, aug 2003.

6. T. de Neergaard, M. Sundwall, and P. Nordenfelt. High-sensitivity assessment of phagocytosis by persistent association-based normalization. BioRxiv, nov 2019.

7. B. Efron. Bootstrap methods: Another look at the jackknife. Ann. Statist., 7(1):1–26, jan 1979.

8. J. J. Ferretti, W. M. McShan, D. Ajdic, D. J. Savic, G. Savic, K. Lyon, C. Primeaux, S. Sezate, A. N. Suvorov, S. Kenton, H. S. Lai, S. P. Lin, Y. Qian, H. G. Jia, F. Z. Najar, Q. Ren, H. Zhu, L. Song, J. White, X. Yuan, S. W. Clifton, B. A. Roe, and R. McLaughlin. Complete genome sequence of an m1 strain of streptococcus pyogenes. Proc Natl Acad Sci USA, 98(8):4658–4663, apr 2001.

9. V. A. Fischetti. M protein and other surface proteins on streptococci. In J. J. Ferretti, D. L. Stevens, and V. A. Fischetti, editors, Streptococcus pyogenes : Basic Biology to Clinical Manifestations. University of Oklahoma Health Sciences Center, Oklahoma City (OK), 2016.

10. A. Forsgren and J. Sjöquist. "protein a" from s. aureus. i. pseudo-immune reaction with human gamma-globulin. J Immunol, 97(6):822–827, dec 1966.

11. I.-M. Frick, P. Åkesson, J. Cooney, U. Sjöbring, K.-H. Schmidt, H. Gomi, S. Hattori, C. Tagawa, F. Kishimoto, and L. Björck. Protein h a surface protein of streptococcus pyogenes with separate binding sites for lgG and albumin. Mol Microbiol, 12(1):143–151, apr 1994.

12. R. W. Glaser. Antigen-antibody binding and mass transport by convection and diffusion to a surface: a two-dimensional computer model of binding and dissociation kinetics. Anal Biochem, 213(1):152–161, aug 1993.

13. S. S. Kadri, B. J. Swihart, S. L. Bonne, S. F. Hohmann, L. V. Hennessy, P. Louras, H. L. Evans, C. Rhee, A. F. Suffredini, D. C. Hooper, D. A. Follmann, E. M. Bulger, and R. L. Danner. Impact of intravenous immunoglobulin on survival in necrotizing fasciitis with vasopressor-dependent shock: A propensity score-matched analysis from 130 US hospitals. Clin Infect Dis, 64(7):877–885, apr 2017.

14. B. M. Kihlberg, M. Collin, A. Olsén, and L. Björck. Protein h, an antiphagocytic surface protein in streptococcus pyogenes. Infect Immun, 67(4):1708–1714, apr 1999.

15. C. Koch, A. Hecker, V. Grau, W. Padberg, M. Wolff, and M. Henrich. Intravenous immunoglobulin in necrotizing fasciitis - a case report and review of recent literature. Ann Med Surg (Lond), 4(3):260–263, sep 2015.

16. J. P. Landry, Y. Ke, G.-L. Yu, and X. D. Zhu. Measuring affinity constants of 1450 monoclonal antibodies to peptide targets with a microarray-based label-free assay platform. J Immunol Methods, 417:86–96, feb 2015.

17. J. J. Lavinder, Y. Wine, C. Giesecke, G. C. Ippolito, A. P. Horton, O. I. Lungu, K. H. Hoi, B. J. DeKosky, E. M. Murrin, M. M. Wirth, A. D. Ellington, T. Dörner, E. M. Marcotte, D. R. Boutz, and G. Georgiou. Identification and characterization of the constituent human serum antibodies elicited by vaccination. Proc Natl Acad Sci USA, 111(6):2259–2264, feb 2014.

18. P. Macheboeuf, C. Buffalo, C.-y. Fu, A. S. Zinkernagel, J. N. Cole, J. E. Johnson, V. Nizet, and P. Ghosh. Streptococcal m1 protein constructs a pathological host fibrinogen network. Nature, 472(7341):64–68, apr 2011.

19. A. J. Martín-Galiano and M. J. McConnell. Using omics technologies and systems biology to identify epitope targets for the development of monoclonal antibodies against antibiotic-resistant bacteria. Front Immunol, 10:2841, dec 2019.

20. M. P. Motley, K. Banerjee, and B. C. Fries. Monoclonal antibody-based therapies for bacterial infections. Curr Opin Infect Dis, 32(3):210–216, 2019.

21. A. N. Nilsson, G. Emilsson, L. K. Nyberg, C. Noble, L. S. Stadler, J. Fritzsche, E. R. B. Moore, J. O. Tegenfeldt, T. Ambjörnsson, and F. Westerlund. Competitive binding-based optical DNA mapping for fast identification of bacteria–multi-ligand transfer matrix theory and experimental applications on escherichia coli. Nucleic Acids Res, 42(15):e118, sep 2014.

22. P. Nordenfelt, S. Waldemarson, A. Linder, M. Mörgelin, C. Karlsson, J. Malmström, and L. Björck. Antibody orientation at bacterial surfaces is related to invasive infection. J Exp Med, 209(13):2367–2381, dec 2012.

23. M. Pedotti, L. Simonelli, E. Livoti, and L. Varani. Computational docking of antibody-antigen complexes, opportunities and pitfalls illustrated by influenza hemagglutinin. Int J Mol Sci, 12(1):226–251, jan 2011.

24. M. I. J. Raybould, W. K. Wong, and C. M. Deane. Antibody–antigen complex modelling in the era of immunoglobulin repertoire sequencing. Mol. Syst. Des. Eng., 4(4):679–688, 2019.

25. V. B. Teif. General transfer matrix formalism to calculate DNA-protein-drug binding in gene regulation: application to OR operator of phage lambda. Nucleic Acids Res, 35(11):e80, may 2007.

26. V. B. Teif and K. Rippe. Calculating transcription factor binding maps for chromatin. Brief Bioinformatics, 13(2):187–201, mar 2012.

27. G. Tibbling, H. Link, and S. Ohman. Principles of albumin and IgG analyses in neurological disorders. i. establishment of reference values. Scand J Clin Lab Invest, 37(5):385–390, sep 1977.

28. U. von Pawel-Rammingen, B. P. Johansson, and L. Björck. IdeS, a novel streptococcal cysteine proteinase with unique specificity for immunoglobulin g. EMBO J, 21(7):1607–1615, apr 2002.

29. P. Åkesson, J. Cooney, F. Kishimoto, and L. Björck. Protein h - a novel IgG binding bacterial protein. Mol Immunol, 27(6):523–531, jun 1990.

30. P. Åkesson, K. H. Schmidt, J. Cooney, and L. Björck. M1 protein and protein h: IgGFc- and albumin-binding streptococcal surface proteins encoded by adjacent genes. Biochem J, 300 (Pt 3):877–886, jun 1994.

